# Expanding The Algal Hydrogen Toolbox: A Non-GMO Platform Reveals Multiple Physiological Routes To Sustained Hydrogen Production Across Microalgae

**DOI:** 10.64898/2026.06.25.734516

**Authors:** Tamar Elman, Roni Amit, Jonathan Tirnover, Andrei Makhon, Jade Rachel Marcus, Sara Jaehnert, Michal Breker, Iftach Yacoby

## Abstract

Sustainable hydrogen production from microalgae remains limited by intrinsic physiological constraints and the need to preserve biomass value for food and feed applications. Transgenic approaches to overcome these limitations were proven successful, yet result in genetically modified (GMO) strains that face major regulatory and deployment barriers. Here, we present a non-GMO experimental platform that enables systematic isolation of hydrogen-producing phenotypes through high-throughput UV mutagenesis pipline coupled with targeted physiological screening. Applying this approach across phylogenetically distinct algal species, including the industrial strain *Chlorella vulgaris* and the extremophile *Chlorella ohadii*, we achieve high discovery efficiency, recovering 0.4-0.6% validated hydrogen-producing mutants and achieving 6.7-25% validation rates among screen-positive candidates, indicating strong enrichment at the primary screening stage. We show that sustained hydrogen production represents a physiologically accessible state emerging across diverse genetic backgrounds. This state is consistently associated with reorganization of photosynthetic electron partitioning, yet arises through multiple distinct configurations that differentially balance hydrogen production, oxygen metabolism, and carbon fixation. This framework provides a scalable route to identify hydrogen-producing strains in industrially relevant algae without introducing foreign DNA and expands the accessible design space for photobiological hydrogen production.

## Introduction

Current efforts in photobiological hydrogen (H_2_) production in microalgae have largely focused on overcoming intrinsic physiological limitations that restrict sustained hydrogen evolution, including the oxygen (O_2_) sensitivity of hydrogenases and competition for reducing power with carbon fixation ^1–7^. Strategies to address these limitations generally fall into two categories. One approach manipulates external cultivation parameters, including light regimes, medium composition, and metabolic supplementation, to shift cellular physiology toward hydrogen production ^8–15^. The other, targets the biological components of the system, modifying the cellular machinery that governs electron flow and hydrogen metabolism ^16–26^. While these approaches have provided important mechanistic insights and extended hydrogen evolution beyond transient states, they largely generate transgenic strains and remain restricted to genetically tractable systems. At the same time, they do not provide a framework for systematically identifying hydrogen-producing states across diverse algal species. Critically, neither approach is designed to address the requirements for industrial deployment. The deployment of photobiological hydrogen systems is ultimately constrained not only by physiology but also by their ability to operate within economically viable and industrially compatible frameworks ^27–29^. This limitation becomes particularly acute at industrial scale, where hydrogen is a low-margin commodity (≈ USD 2 kg^−1^) ^30,31^. As a result, photobiological systems based solely on hydrogen output are unlikely to be economically viable without additional value streams. Microalgae offer a distinct advantage in this context: alongside hydrogen, they generate biomass with established value in food and feed markets ^32–34^. Co-valorization of biomass is therefore not optional but a central design constraint that directly shapes strain selection and cultivation strategies ^28^.

The genetic status of production strains becomes a critical factor under this framework. Regulatory definitions in major markets restrict the use of genetically modified organisms in food and feed applications, thereby limiting the deployment of transgenic algal strains ^35,36^. These limitations motivate the development of non-transgenic approaches for strain improvement that remain compatible with downstream applications ^37–39^. Non-transgenic mutagenesis approaches can generate heritable variation within native genetic backgrounds, but their effectiveness depends on the ability to identify relevant phenotypes at scale. In microalgae, UV mutagenesis has been extensively applied to improve traits such as growth and lipid accumulation, often in combination with high-throughput screening approaches ^40–44^. However, its application to hydrogen production has remained limited, largely due to the lack of scalable and selective screening strategies capable of resolving relevant physiological phenotypes ^45,46^. For example, Bayro-Kaiser and Nelson demonstrated that UV mutagenesis can generate hydrogen-relevant phenotypes in *Chlamydomonas reinhardtii*, but the reliance on indirect selection and low recovery rates (∼0.033%) highlights the difficulty of identifying such traits in a scalable manner ^46^.

Here, we address this gap by establishing a scalable, non-GMO discovery platform designed to identify hydrogen-producing phenotypes across phylogenetically diverse algal species (Fig. 1). The platform combines UV mutagenesis with targeted physiological screening to systematically recover mutants capable of sustained hydrogen production from a baseline of transient activity. Rather than optimizing a single engineered pathway, this approach enables exploration of multiple physiological configurations that support hydrogen production. A central feature of this workflow is a high-light selection step (∼900 µmol photons m^−2^ s^−1^), which imposes strong excitation pressure and enables rapid filtration of mutants unable to appropriately manage photosynthetic electron flux. This selection enriches for strains with altered electron allocation, a physiological regime associated with hydrogen production.

**Figure 1.**
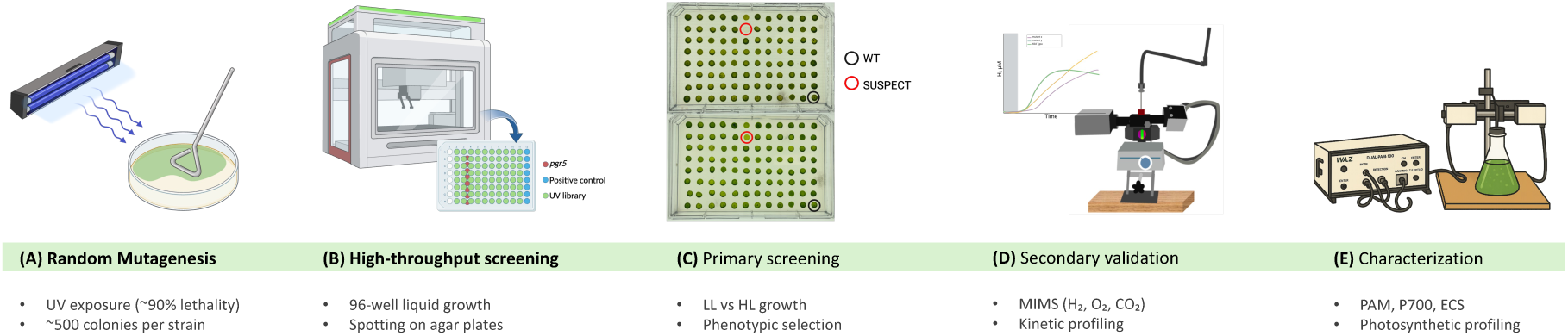
High-throughput UV mutagenesis and screening workflow. (**A**) Random mutagenesis was performed using UV irradiation. Exposure times were calibrated for each algal strain to achieve ∼90% lethality, generating mutant libraries of approximately 500 colonies per strain. (**B**) Mutant libraries were grown in 96-well plates under low-light conditions to accumulate biomass and subsequently arrayed on agar plates alongside wild-type controls. (**C**) Primary phenotypic screening was based on differential growth under low-light (∼30 μmol photons m⁻² s⁻¹) versus high-light (∼900 μmol photons m⁻² s⁻¹) conditions to identify candidate mutants. (**D**) Selected candidates were subjected to secondary validation using membrane inlet mass spectrometry (MIMS) to quantify hydrogen production and associated gas exchange dynamics (H₂, O₂, and CO₂). (**E**) Validated mutants were further characterized using complementary physiological measurements, including chlorophyll fluorescence (PAM), P700 redox kinetics, and electrochromic shift (ECS), to resolve photosynthetic and metabolic properties associated with sustained hydrogen production.

Using this approach, we achieve substantially improved discovery efficiency and screening precision relative to previous methods, with true-positive rates of ∼0.4–0.6% across multiple species, including *Chlamydomonas reinhardtii*, *Chlorella vulgaris*, and *Chlorella ohadii*. Importantly, this improvement reflects not only higher recovery from the initial libraries, but also strong enrichment at the high-light screening stage, with 6.7–25% of screen-positive candidates subsequently validated as true H_2_-producing mutants. This cross-species applicability enables diversification of hydrogen-producing strains and supports alignment with different biomass markets. Beyond strain identification, the platform provides a comparative physiological framework to resolve how distinct photosynthetic systems reallocate electron flow to sustain hydrogen production. Together, these results establish a scalable, non-GMO discovery framework that enables systematic identification of hydrogen-producing states across diverse algal systems and expands the range of accessible physiological solutions for photobiological hydrogen production.

## 3. Results

### 3.1 Establishment of a high-throughput UV-mutagenesis pipeline

To enable systematic identification of hydrogen-producing phenotypes, we first established a UV-mutagenesis pipeline optimized for downstream screening. Exposure times were calibrated separately for each algal species to achieve ∼90% mortality (∼10% survival), ensuring a substantial mutational load while maintaining recoverable colony numbers, thereby maximizing phenotypic diversity and preserving a manageable library size. Despite applying a consistent survival criterion, required exposure times varied markedly among species, highlighting substantial differences in intrinsic UV tolerance across phylogenetically distinct algae. *Chlorella ohadii* required the longest treatment (2:45 min), *Chlamydomonas reinhardtii* strains CC-125 and T222 required 1:10 min and 0:45 min, respectively, whereas *Chlorella vulgaris* was most sensitive, requiring only 0:35 s.

Following mutagenesis optimization, we defined the starting library size for screening. We selected an initial scale of 500 colonies per strain. This size balances phenotypic diversity with practical feasibility, chosen to enable complete downstream validation of all candidates while maintaining sufficient diversity to recover hydrogen-producing phenotypes. Importantly, this scale was sufficient to recover multiple independent hydrogen-producing mutants across all tested species, demonstrating that the pipeline achieves meaningful enrichment without requiring large libraries. The workflow remains inherently scalable and can be expanded to larger libraries under automated conditions as the screening logic is independent of library size.

### 3.2 Primary screen for hydrogen producers

Efficient identification of hydrogen-producing mutants presents a major bottleneck, as reliable quantification requires mass spectrometry and is not compatible with high-throughput screening. We therefore implemented a rapid primary screen designed to enrich for candidates with altered photosynthetic regulation prior to quantitative validation.

As a benchmark, we used the well-characterized *pgr5* mutant, which sustains hydrogen production under ambient conditions but exhibits pronounced sensitivity to high light due to impaired cyclic electron flow around photosystem I (PSI), resulting in altered redox balance and reduced photoprotective capacity ^47^. We reasoned that additional mutants with defective photosynthetic regulation, and consequently high-light sensitivity, may present with related physiological states that favor hydrogen production. High-light sensitivity was therefore used as a phenotypic proxy to enrich for candidates with altered electron partitioning, rather than as a direct predictor of hydrogen production.

For each library, 500 colonies were collected manually or using an automated picker and grown in liquid TAP medium in 96-well plates, with the wild type included as an internal control. After biomass accumulation under low light, cultures were arrayed onto TAP agar in duplicate (Figure 2). One plate was maintained under low light (∼30 μmol photons m^−2^ s^−1^) as a growth control, while the second was exposed to high light (∼900 μmol photons m^−2^ s^−1^) to impose photoinhibitory stress. For *Chlorella ohadii*, an additional plate was exposed to natural sunlight to apply an extreme high-light challenge. Screening was performed during peak summer irradiance (up to ∼3500 μmol photons m^−2^ s^−1^). Under these conditions, *C. ohadii* was uniquely capable of sustained growth, consistent with its exceptional high-light tolerance ^48^, thereby providing a stringent benchmark for mutant phenotyping.

**Figure 2.**
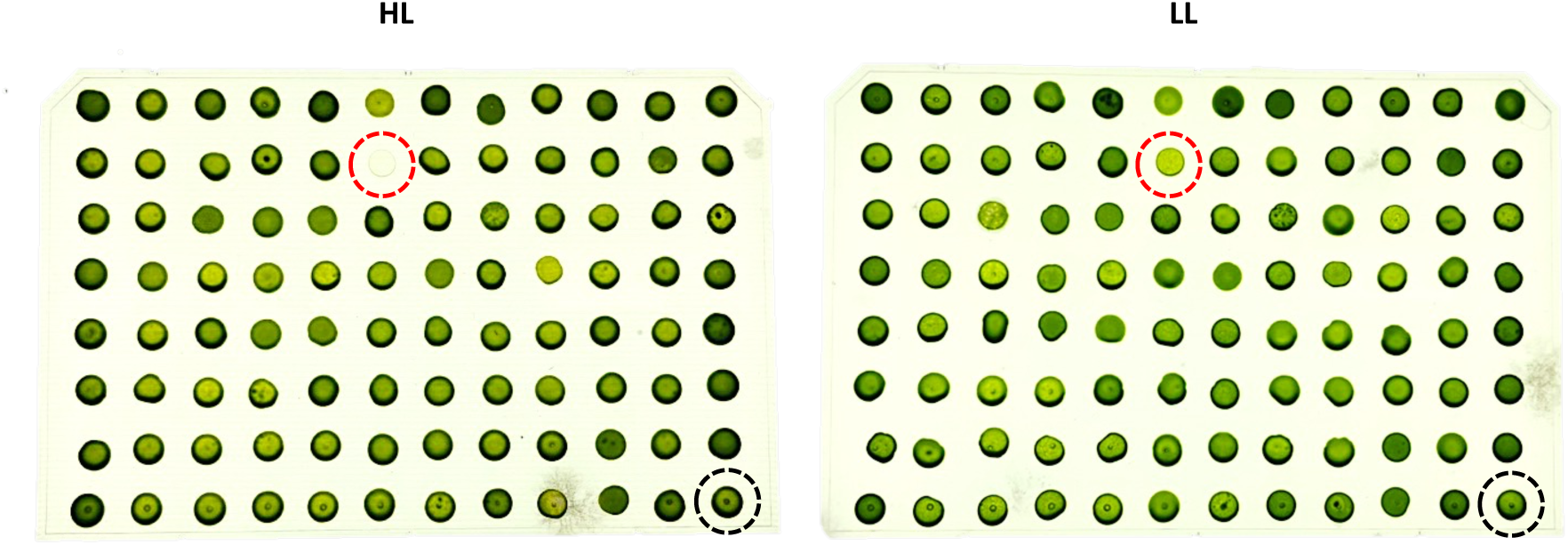
High-throughput high-light screening for H_2_-producing candidates. Colonies from 96-well cultures were spotted onto duplicate TAP agar plates and incubated under low light (LL; 30 µmol photons m^−2^ s^−1^; right plate) or high white light (HL; 900 µmol photons m^−2^ s^−1^; left plate). For *Chlorella ohadii*, an additional plate was exposed to direct sunlight. Growth under LL confirms colony viability, whereas failure or reduced growth under HL/sun identifies high-light–sensitive candidates selected for downstream H_2_ validation. The wild-type control is indicated by a black dashed circle; a representative mutant identified in the screen and selected for follow-up validation is marked by a red dashed circle.

After 5 days, colonies that failed under high light yet grew normally under control conditions were classified as screen-positive candidates. To minimize false positives, these candidates were re-screened in an independent round under identical conditions. Only colonies that reproducibly displayed the phenotype were retained for downstream hydrogen validation.

### 3.3 Identification of true positives using membrane inlet mass spectrometry

Following the primary screen, hydrogen production was validated using membrane inlet mass spectrometry (MIMS), as high-light sensitivity alone is not sufficient to establish hydrogen-producing capacity. Individual screen-positive candidates were analyzed by MIMS under anaerobic conditions. Although gas chromatography can be used to quantify hydrogen and represents a more widely accessible method, MIMS provides real-time, high-resolution monitoring of H_2_, O_2_, and CO_2_ simultaneously, allowing direct assessment of the gas-exchange dynamics underlying sustained hydrogen production. In an initial survey, only a subset of candidates displayed clear and reproducible H_2_ evolution. These mutants were subsequently validated in biological replicates (Figure 3).

**Figure 3.**
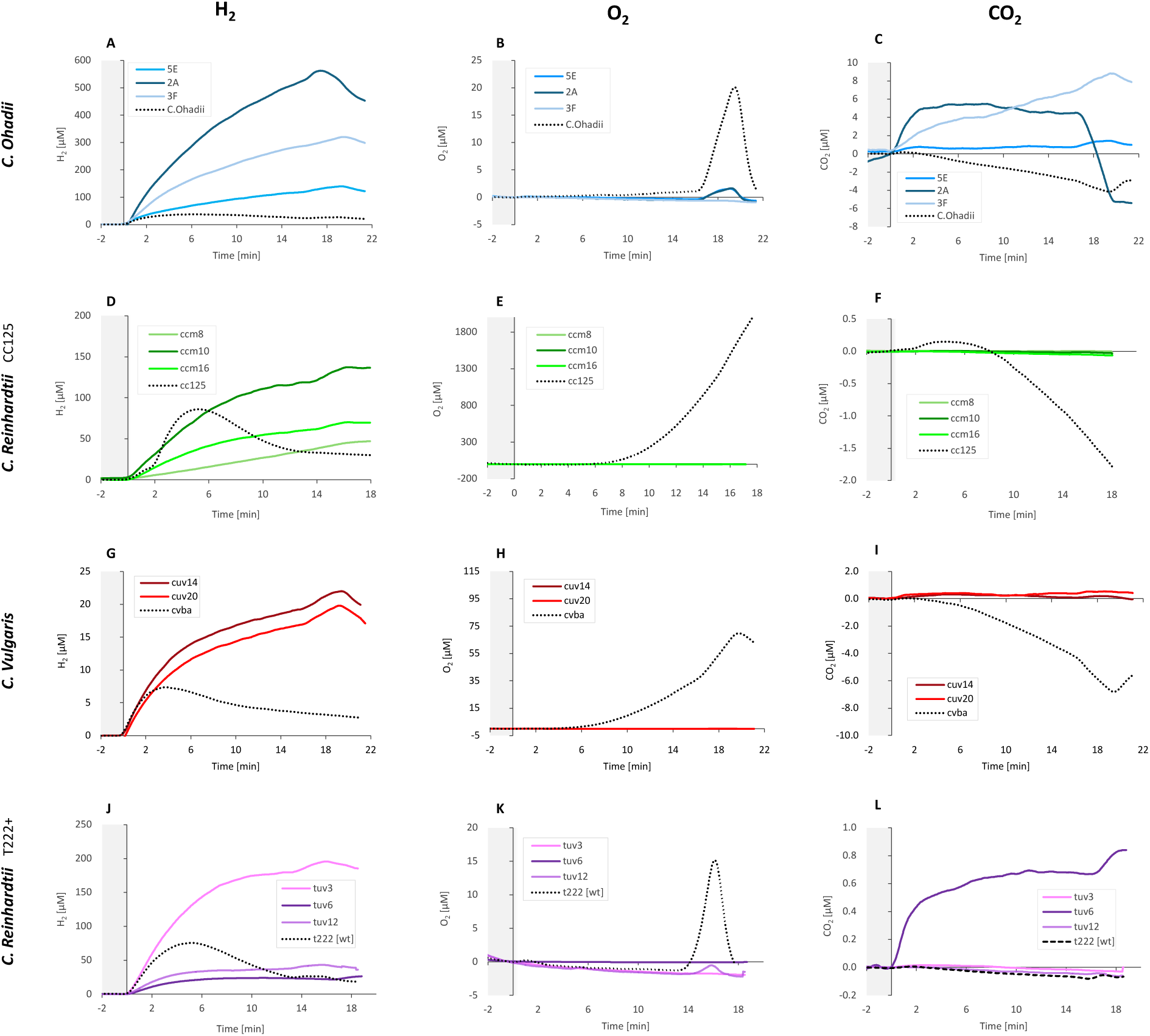
Identification of true positive hydrogen-producing mutants using MIMS analysis. Colonies that passed the primary light-based screen were analyzed for gas-exchange dynamics using MIMS. Cultures adjusted to ∼20 μg Chl mL^−1^ were dark-incubated for 2 h, followed by illumination at 370 μmol photons m^−2^ s^−1^ for 16 min, a 3-min pulse at saturating light (2,500 μmol photons m^−2^ s^−1^), and a final 2-min dark period. Each row represents one wild-type algal species (dashed black curves) and its corresponding mutants (solid-colored curves). (A, D, G, J) Hydrogen (H_2_), (B, E, H, K) oxygen (O_2_), and (C, F, I, L) carbon dioxide (CO_2_) exchange. Values represent mean values (n ≥ 3).

The MIMS assay consisted of a two-hour dark incubation to deplete residual oxygen, followed by 16 minutes of actinic light illumination at 370 μmol photons m^−2^ s^−1^ and a final 3-minute pulse at saturating light (∼2500 μmol photons m^−2^ s^−1^). Wild-type strains displayed the expected transient H_2_ burst, which peaked early and collapsed within 2–5 minutes of illumination (Figure 3A, D, G, J). In contrast, selected mutants maintained hydrogen evolution beyond this initial transient phase, resulting in sustained H_2_ production throughout the illumination period.

Across species, two to three mutants per background exhibited sustained hydrogen production under these conditions. In *Chlorella ohadii* (Figure 3A–C), mutants 2A, 5E, and 3F displayed continuous H_2_ evolution at rates exceeding those of the wild type. Similarly, in *Chlorella vulgaris* (Figure 3G–I), mutants cuv14 and cuv20 produced hydrogen more efficiently and for longer durations than the parental strain. In *Chlamydomonas reinhardtii*, greater variability was observed. In both the CC125 background (Figure 3D–F; mutants ccm8, ccm10, ccm16) and the T222 background (Figure 3J–L; mutants tuv3, tuv6, tuv12), several strains sustained hydrogen evolution throughout the illumination period. Although peak rates were often lower than the transient maxima observed in the respective wild types, prolonged production resulted in increased cumulative H_2_ output in selected mutants, as seen in 3D.

Across all validated strains, hydrogen production was accompanied by alterations in O_2_ and CO_2_ exchange relative to wild-type controls. O_2_ levels remained negligibly low during illumination, although the extent varied between strains (Figure 3B, E, H, K). In parallel, CO_2_ depletion was slower or incomplete relative to wild type (Figure 3C, F, I, L), indicating attenuated carbon fixation under these conditions.

Together, these gas-exchange patterns define a consistent physiological signature associated with sustained hydrogen production.

Importantly, sustained hydrogen production was not associated with severe impairment of biomass accumulation. Across all genetic backgrounds, mutants maintained growth profiles broadly comparable to their parental lines under standard cultivation conditions, with only moderate reductions in final density observed in specific cases (Supplementary Fig. 1). This decoupling from major growth penalties distinguishes these strains from classical stress-induced hydrogen production systems (such as sulfur-deprivation method^49^) and supports their relevance for cultivation-based applications.

To quantify the overall performance of the UV-mutagenesis and screening workflow, we summarized the full pipeline, from mutagenesis to confirmed H_2_-positive mutants, in Table 1. Each screen began with a library of 500 colonies per strain, generated under species-specific UV conditions calibrated to yield ∼10% survival. Following the high-light growth assay, 2-6% of colonies were classified as screen-positive candidates from the initial library. Subsequent validation by MIMS reduced this fraction to 0.4-0.6% of the original library, with three backgrounds converging at ∼0.6% and Chlorella vulgaris at 0.4%. Importantly, the primary screen exhibited high precision, with 6.7–25% of screen-positive candidates validated as true H_2_-producing mutants, representing a substantial enrichment relative to the initial library. For comparison, a previous UV-mutagenesis effort in Chlamydomonas reinhardtii CC125 reported ∼0.03% true positives from a substantially larger initial library (n = 12,000). In contrast, the present pipeline reproducibly yields a more than 10-fold higher success rate of 0.4–0.6% validated hydrogen-producing mutants across multiple algal species, despite starting from a smaller and manually manageable library size. Together, these results demonstrate that the screening strategy improves both the recovery and the selectivity of hydrogen-producing phenotypes.

**Table 1.** Summary of UV-mutagenesis workflow efficiency across algal species. Approximately 5,000 cells per strain were subjected to UV irradiation under species-specific conditions yielding ∼10% survival and ∼500 colonies (mutant library). Colonies identified as high-light sensitive in the primary screen are reported as “screen-positive candidates,” and mutants validated by MIMS as sustained hydrogen producers are classified as “true positives.” Two complementary performance metrics are shown: (i) overall recovery efficiency, calculated as the fraction of the initial library yielding confirmed hydrogen-producing mutants, and (ii) primary screen precision, defined as the percentage of screen-positive candidates subsequently validated by MIMS as true positives.

|  | +UV | *C. Ohadii* |  | *C. Reinhardtii* (CC125) |  | *C. Vulgaris* |  | *C. Reinhardtii* (T222+) |  |
| --- | --- | --- | --- | --- | --- | --- | --- | --- | --- |
|  |  | Positives suspects | True positives | Positives suspects | True positives | Positives suspects | True positives | Positives suspects | True positives |
| Colonies # | 500 | 15 | 3 | 20 | 3 | 30 | 2 | 12 | 3 |
| % from initial library | 100 % | 3 % | 0.6 % | 4% | 0.6% | 6 % | 0.4 % | 2.4% | 0.6 % |
| Primary screen precision (%) |  |  |  |  |  |  |  |  |  |
|  |  |  | 20 % |  | 15 % |  | 6.7 % |  | 25 % |

### 3.4 Insights into photosynthetic function from physiological assays

Having established sustained hydrogen production across multiple species, we next sought to characterize the physiological basis of this phenotype. We therefore assessed oxygen dynamics, photosynthetic electron partitioning, and thylakoid energetic states in the highest hydrogen-producing mutant from each species to identify physiological adjustments associated with sustained hydrogen evolution.

#### 3.4.1 Oxygen dynamics revealed by PI curves

To examine how O_2_ dynamics are modulated in hydrogen-producing mutants, we quantified PSII-driven O_2_ evolution across increasing irradiances using a fiber-optic O_2_ probe. Dark-adapted cultures were exposed to successive 1-min light-dark cycles from 50 to 2,000 µmol photons m^−2^ s^−1^, generating light-response photoinhibition (PI) curves for each mutant and its corresponding wild type. Across strains, two distinct patterns were observed: (i) sustained PSII-driven O_2_ evolution coupled to increased O_2_ consumption, and (ii) intrinsically reduced PSII output limiting O_2_ production.

In *Chlorella ohadii* 2A (Fig. 4A) and *Chlamydomonas reinhardtii* ccm10 (Fig. 4B), gross O_2_ evolution continued to increase with irradiance and did not exhibit the saturation behavior observed in the respective wild types, indicating that PSII activity remains responsive to increasing light input. However, net O_2_ accumulation was attenuated relative to gross O_2_, particularly at high irradiance, resulting in a clear divergence between the two. This pattern is consistent with enhanced O_2_ consumption downstream of PSII, rather than reduced O_2_ production.

**Figure 4.**
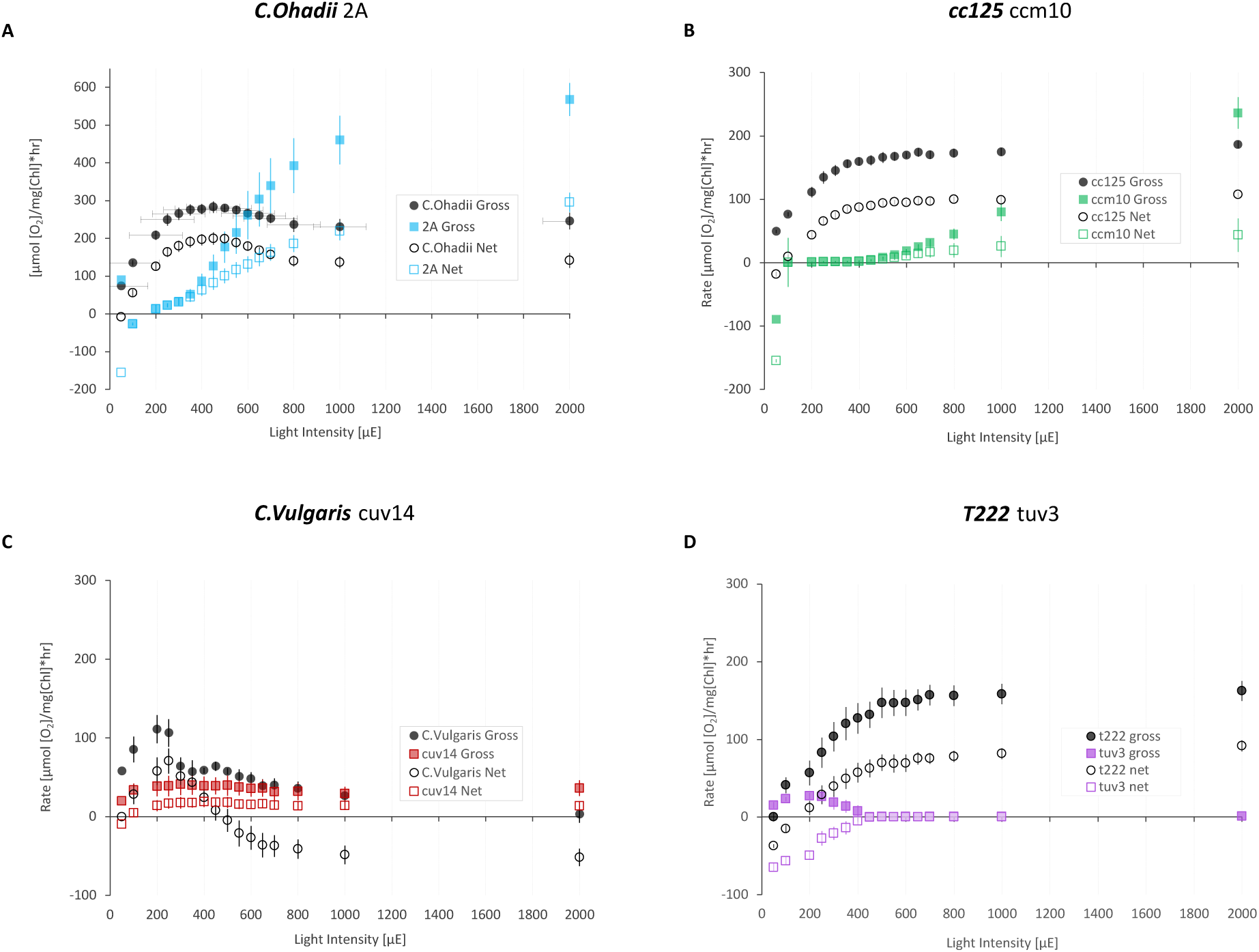
Light response curves of net and gross O_2_ evolution in wild-type and hydrogen-producing mutant strains. Gross and net O_2_ evolution were measured under oxic conditions using a PyroScience FireSting fiber-optic O_2_ probe in cultures adjusted to ∼20 μg Chl mL^−1^. Dissolved O_2_ (μM) was continuously recorded during successive 1-min light-dark cycles under white light at irradiances ranging from 50 to 2000 μmol photons m⁻² s^−1^. Gross O_2_ evolution was calculated by subtracting the preceding dark respiration rate from the light-induced O_2_ signal, while net O_2_ evolution represents the directly measured O_2_ exchange. Each panel corresponds to one algal species, shown as the wild type (black symbols) and its highest H_2_-producing mutant (colored symbols). Filled symbols indicate gross O_2_ evolution; open symbols indicate net O_2_ evolution. (A) *Chlorella ohadii* vs. 2A. (B) *Chlamydomonas reinhardtii* CC125 vs. ccm10. (C) *Chlorella vulgaris* vs. cuv14. (D) *Chlamydomonas reinhardtii* T222 vs. tuv3. Values represent mean ± SE (n ≥ 3).

A contrasting pattern was observed in *Chlorella vulgaris* cuv14 (Fig. 4C) and *C. reinhardtii* tuv3 (Fig. 4D). In these strains, gross O_2_ evolution was reduced across irradiances, indicating intrinsically limited PSII activity. In cuv14, gross O_2_ remained low and plateaued without the decline observed in the wild type, while the separation between gross and net O_2_ remained relatively constant, suggesting a stable balance between production and consumption. In tuv3, gross O_2_ declined at intermediate irradiances, and net O_2_ remained negative at low light, approaching zero at higher irradiance, consistent with a supply-limited regime in which O_2_ production is insufficient to offset consumption.

These results resolve the oxygen phenotype into two distinct physiological regimes: one characterized by sustained PSII-driven O_2_ evolution coupled to increased O_2_ consumption, and the other defined by intrinsically reduced PSII output limiting O_2_ production.

#### 3.4.2 Assessment of PSII activity and energy dissipation by PAM fluorometry

To determine whether the distinct oxygen-response regimes reflect differences in PSII function, we examined PSII photochemistry and energy dissipation using chlorophyll fluorescence measurements conducted in parallel with the MIMS gas-exchange assays (Fig. 5).

**Figure 5.**
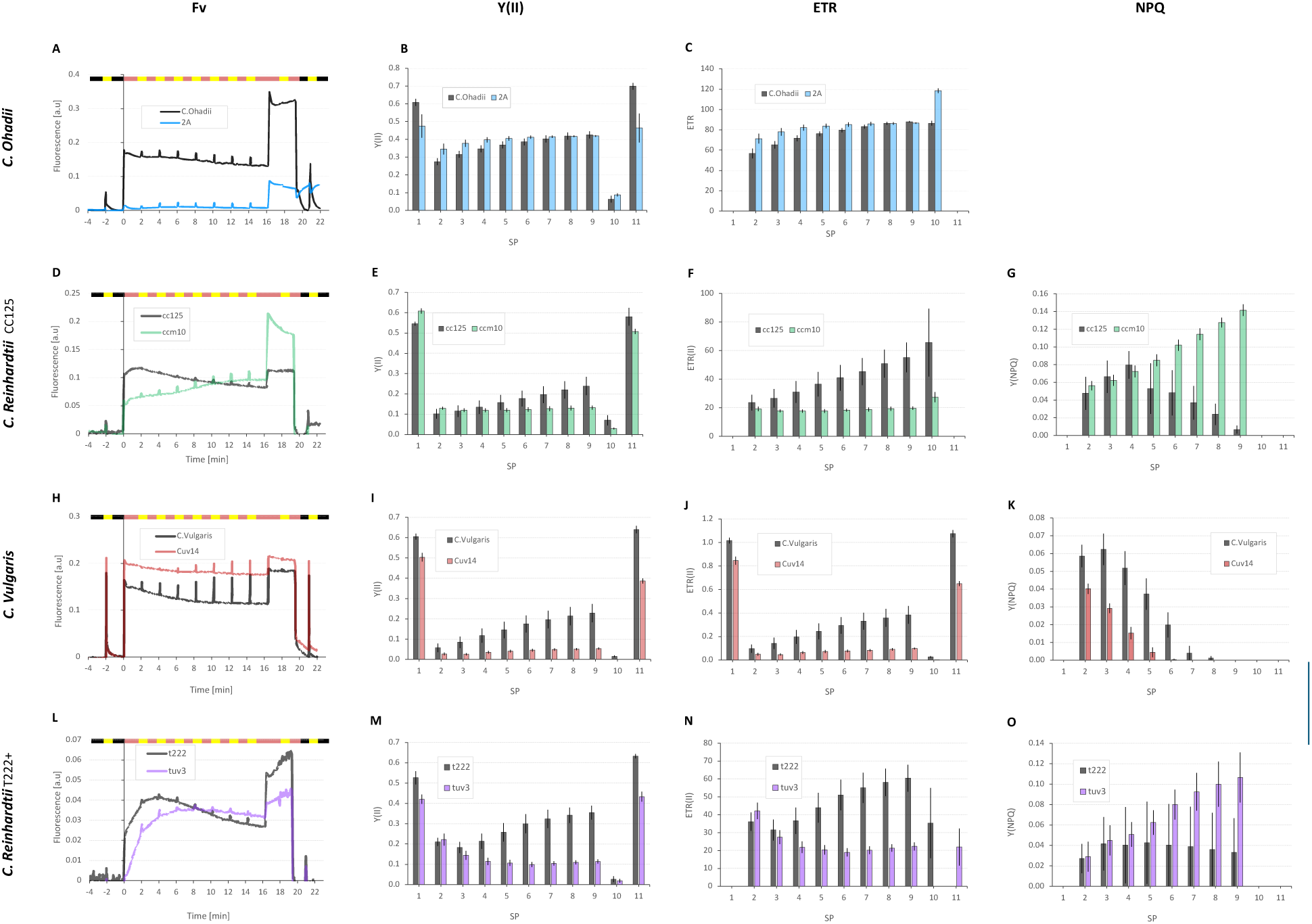
Comparative photosynthetic characterization of wild-type and mutant algal strains by chlorophyll fluorescence. Chlorophyll fluorescence was measured in parallel with MIMS gas exchange (Fig. 2) using a Dual-PAM-100 fluorometer (Walz). Each row represents a different algal species, shown as the wild type (black) and the corresponding hydrogen-producing mutant selected for further analysis. (A, D, H, L) Fluorescence traces showing minimum (F_0_) and maximum (Fm) fluorescence yields measured by saturating pulses (SP, arrows) following dark adaptation (left black bar). During actinic illumination (red bar), fluorescence quenching occurred, and Fm’ was determined by SPs. Additional SPs were applied during the subsequent dark recovery phase (right black bar). Derived parameters are shown in the adjacent panels: (B, E, I, M) effective quantum yield of PSII (ΦPSII), (C, F, J, N) electron transport rate (ETR), and (G, K, O) non-photochemical quenching (NPQ). Data represent mean ± SEM (n ≥ 3).

Wild-type strains displayed canonical fluorescence responses, including a transient peak upon transition to actinic light followed by a gradual decline (Kautsky effect), together with increasing Y(II) and electron transport rate (ETR) and transient NPQ formation, consistent with efficient photochemistry and balanced energy dissipation.

Mutants exhibiting reduced net O_2_ accumulation showed distinct PSII alterations. *Chlorella vulgaris* cuv14 and *Chlamydomonas reinhardtii* tuv3, which display intrinsically low gross O_2_ evolution in the PI curves, showed consistently low and largely unresponsive Y(II) and ETR across irradiances (Fig. 5I–J, M–N), indicating a limited capacity to dynamically activate photochemistry under actinic light. This behavior is consistent with their reduced PSII-driven O_2_ output observed in the PI curves.

NPQ responses varied between these strains (Fig. 5K, O): tuv3 exhibited a light-dependent increase in NPQ, whereas cuv14 retained a qualitatively wild-type-like NPQ pattern but at lower amplitude. In cuv14, the presence of fluorescence decay alongside WT-like NPQ dynamics indicates effective excitation quenching; however, the persistently low Y(II) and ETR suggest that this quenching is not efficiently coupled to photochemistry.

A distinct pattern was observed in *Chlamydomonas reinhardtii* ccm10. Despite exhibiting sustained gross O_2_ evolution under increasing irradiance, ccm10 also displayed reduced Y(II) and ETR (Fig. 5E–F), indicating that its oxygen phenotype does not arise from uniformly enhanced PSII activity across light regimes. This prompted closer examination of the measurement regimes. Importantly, this apparent discrepancy reflects the different light regimes probed by the two measurements. At moderate irradiances (300–400 μmol photons m^−2^ s^−1^), corresponding to the PAM measurements, both gross and net O_2_ evolution in ccm10 remain low and closely overlapping (Fig. 4B), consistent with the low Y(II) and ETR values observed. Only at higher irradiances does PSII activity increase, resulting in the elevated gross O_2_ evolution captured in the PI curves.

Taken together, this pattern suggests a more complex regulation of electron flow, in which PSII output is constrained at moderate light but becomes engaged at higher irradiance, while downstream processes contribute to limiting net O_2_ accumulation.

In contrast, *Chlorella ohadii* mutant 2A represents the regime of sustained PSII activity. This strain closely matched its wild type across all measured fluorescence parameters, including Y(II) and ETR dynamics (Fig. 5B–C), indicating preserved PSII photochemistry despite reduced net O_2_ accumulation in the PI curves. This supports the interpretation that O_2_ limitation in this strain arises from enhanced downstream consumption rather than reduced PSII activity.

Although absolute fluorescence amplitudes (F_0_ and Fm) were lower than in the other species (Fig. 5A), this reflects the intrinsic high-light-adapted physiology of *C. ohadii* rather than mutant-specific effects. NPQ analysis was not included for this strain due to its minimal NPQ response under these conditions^48^ (Fig. 5D).

#### 3.4.3 PSI redox partitioning and thylakoid energetic states in H_2_-producing mutants

To further resolve how electron flow is partitioned downstream of PSII, we analyzed thylakoid energetic states and PSI redox dynamics. Electrochromic shift (ECS) measurements were first used to assess thylakoid membrane potential dynamics (Δψ) (Fig. 6). In parallel, PSI activity was quantified using P700 oxidation measurements, allowing decomposition of the oxidizable PSI pool into photochemically active PSI (Y(I)), acceptor-side limitation (YNA), and donor-side limitation (YND) (Fig. 7). Representative P700 oxidation kinetics underlying these quantifications are shown in Supplementary Fig. S2.

**Figure 6.**
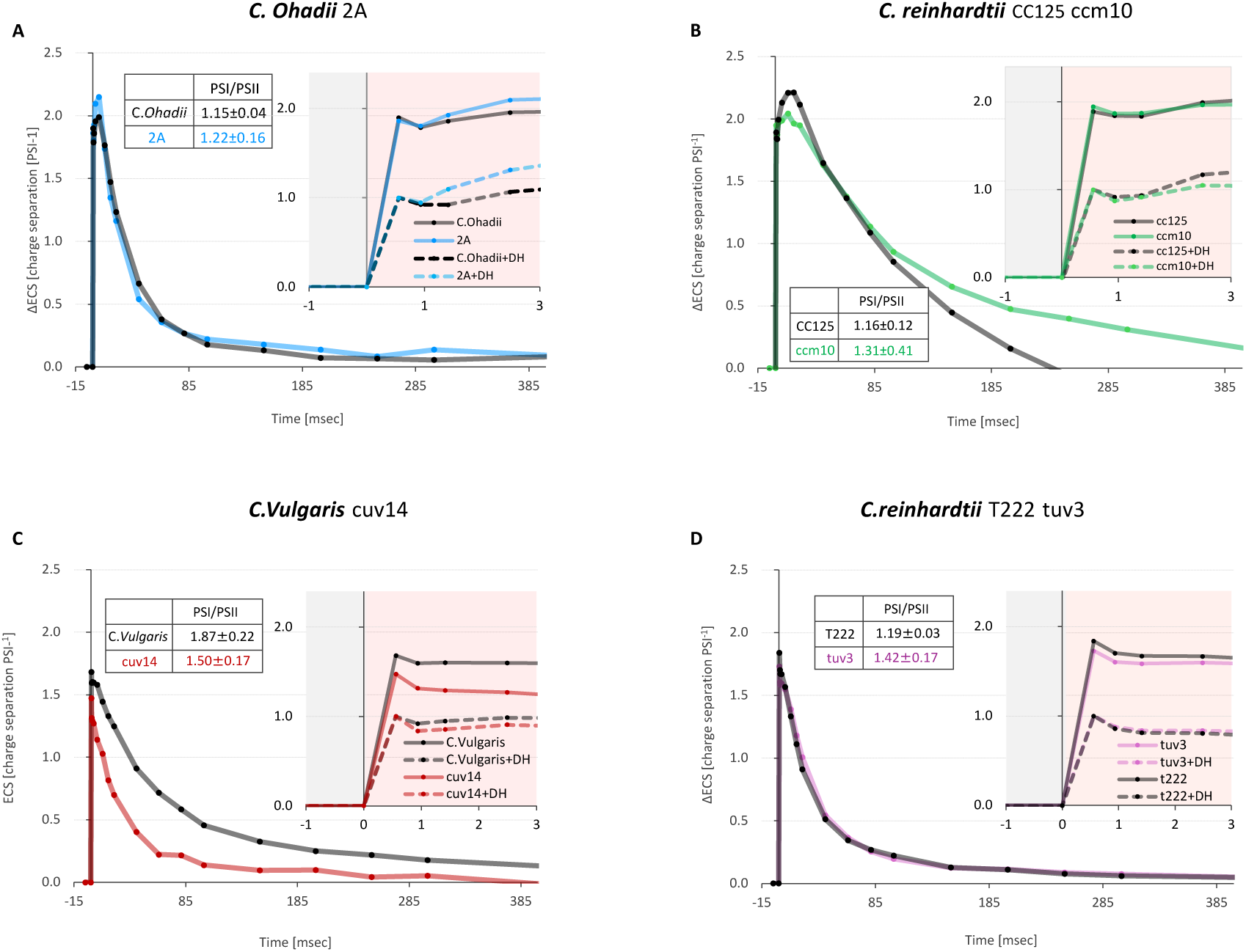
ECS measurements of PSI/PSII ratio and membrane potential relaxation in wild-type and mutant algal strains. Electrochromic shift (ECS) kinetics were recorded under oxic conditions using a Joliot-type spectrophotometer (JTS-150) on cells adjusted to ∼20 μg Chl mL^−1^ to evaluate charge separation, PSI/PSII ratios, and relaxation of the thylakoid membrane potential (Δψ). Each panel corresponds to a different algal species, with wild type (black traces) compared to its highest H₂-producing mutant (colored traces): (A) *Chlorella ohadii* vs. 2A, (B) *Chlamydomonas reinhardtii* CC125 vs. ccm10, (C) *Chlorella vulgaris* vs. cuv14, and (D) *Chlamydomonas reinhardtii* T222 vs. tuv3. Insets show magnified views of the rapid induction phase and a summary table of PSI/PSII ratios (mean ± SE, n ≥ 3), calculated from measurements performed in the presence of 10 μM DCMU and 1 mM hydroxylamine (+DH traces) to block PSII.

**Figure 7.**
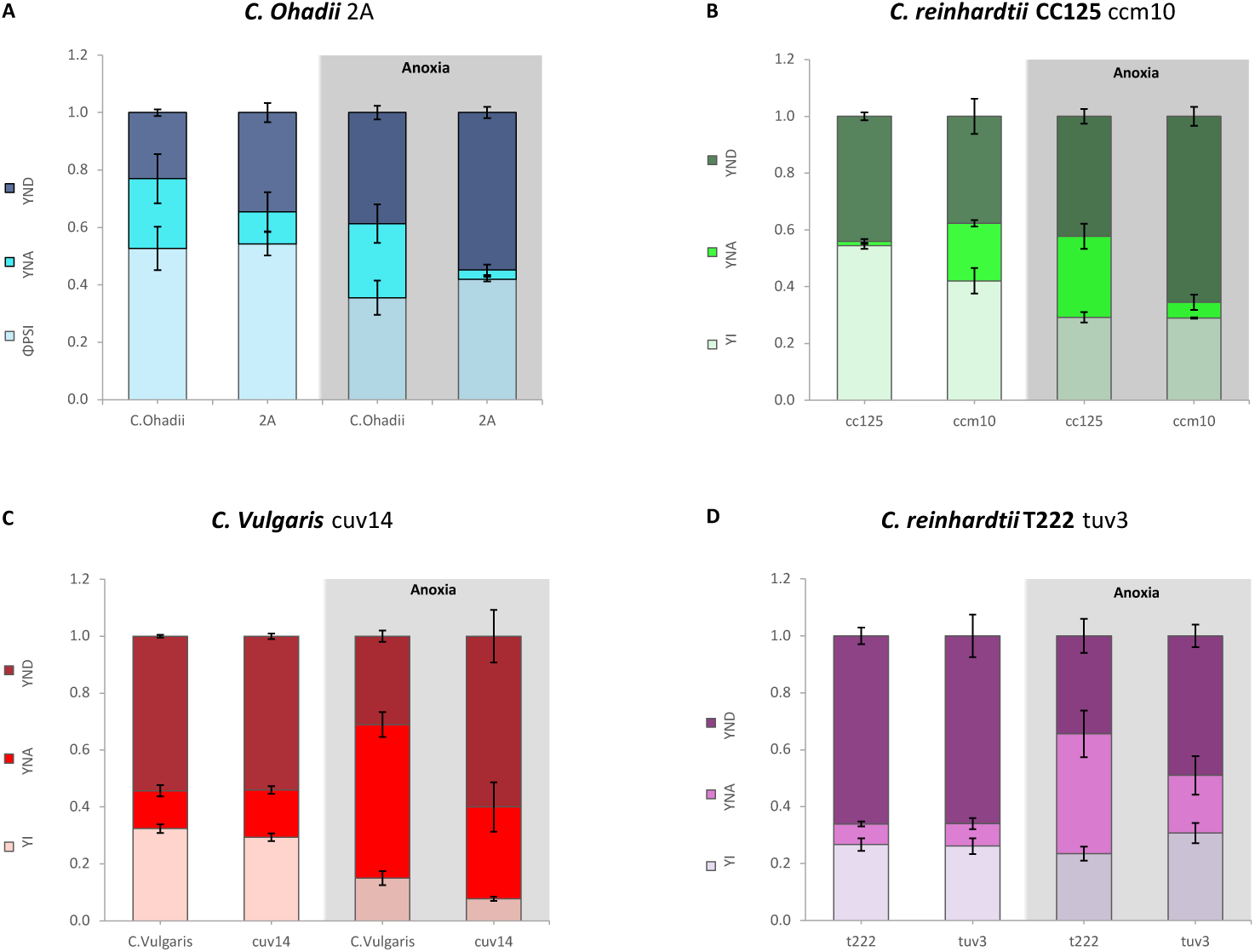
PSI redox partitioning in hydrogen-producing mutants. PSI redox partitioning derived from P700 oxidation measurements. Stacked bars represent the relative fractions of total P700 assigned to photochemically active PSI (Y(I)), acceptor-side limitation (YNA), and donor-side limitation (YND). Each panel corresponds to a different species, shown as wild type and its highest hydrogen-producing mutant. For each strain, bars compare aerobic conditions (left) with samples subjected to 2 h dark anaerobic incubation (right, grey background). Values represent means ± SE (n ≥ 3 biological replicates). Quantification is based on P700 oxidation kinetics shown in Supplementary Fig. S2.

ECS measurements revealed divergence in thylakoid energetic states between mutants (Fig. 6), providing insight into how proton motive force and membrane potential dynamics are regulated across strains. While *Chlorella ohadii* 2A and *Chlamydomonas reinhardtii tuv3* displayed ECS kinetics broadly similar to their respective wild types under aerobic conditions, other mutants exhibited clear deviations in both amplitude and relaxation dynamics. In *Chlamydomonas reinhardtii ccm10*, Δψ relaxation was slower, suggesting prolonged retention of membrane potential and altered proton flux. In contrast, *Chlorella vulgaris cuv14* showed reduced ECS amplitude and faster relaxation, consistent with limited Δψ buildup and increased proton conductivity.

Across all strains, PSI redox partitioning shifted in a condition-dependent manner (Fig. 7). Under anoxic conditions, all mutants exhibited a relative increase in donor-side limitation (YND) compared to their respective wild types, indicating restricted electron delivery to PSI. This convergence suggests that, despite diverse upstream configurations, sustained hydrogen production is associated with a common redox state in which PSI becomes increasingly donor limited.

Despite this shared outcome, the underlying partitioning patterns differed markedly between strains. In *Chlorella ohadii* mutant 2A (Fig. 7A), YNA was consistently lower than in the wild type under both aerobic and anoxic conditions, indicating reduced acceptor-side limitation and supporting enhanced downstream electron withdrawal. This behavior is consistent with the preserved PSII activity and elevated O_2_ consumption observed in this strain. In *Chlamydomonas reinhardtii ccm10* (Fig. 7B), PSI partitioning exhibited a pronounced shift between conditions. Under aerobic conditions, YNA was elevated relative to the wild type, indicating acceptor-side limitation. Under anoxia, however, YND became dominant, reflecting restricted electron supply. This transition indicates that PSI activity in this mutant is not fixed but depends strongly on the redox environment, consistent with the complex regulation inferred from the O_2_ and PAM measurements. A different configuration was observed in *Chlorella vulgaris cuv14* (Fig. 7C), where PSI partitioning under anoxia was strongly shifted toward donor-side limitation, accompanied by a marked reduction in Y(I). This behavior is consistent with the low PSII activity and supply-limited regime identified in the PI curves and PAM measurements. Similarly, in *Chlamydomonas reinhardtii tuv3* (Fig. 7D), PSI partitioning under aerobic conditions resembled the wild type, whereas under anoxia a clear shift toward donor-side limitation was observed. This condition-dependent redistribution supports a model in which electron flow becomes constrained primarily at the level of supply during hydrogen-producing conditions.

Together, these results indicate that sustained hydrogen production is not associated with a single mechanistic pathway but rather emerges from multiple physiological routes. While PSI redox partitioning converges toward donor-side limitation under anoxia, thylakoid energetic states remain highly variable, indicating that distinct energetic configurations can support similar downstream redox outcomes.

### 3.5 Whole-genome sequencing and prioritization of candidate loci in the *Chlorella ohadii* mutant 2A

To translate the physiological phenotypes identified above into genetic hypotheses, we performed whole-genome sequencing of the most robust H_2_-producing mutant, *Chlorella ohadii* 2A, as a final step in the discovery pipeline. Beyond identifying candidate mutations, this analysis demonstrates how strains emerging from the screening platform can be advanced toward mechanistic interrogation and targeted follow-up. Rather than establishing direct causality, the approach was designed to prioritize candidate *loci* associated with the observed phenotype.

To distinguish mutagenesis-derived variants from background polymorphisms, the parental wild-type strain maintained in our laboratory was sequenced in parallel. Both genomes were independently compared to the publicly available *Chlorella ohadii* reference genome, and only variants uniquely present in the mutant and absent from the laboratory wild type were retained for further analysis. This filtering strategy removes strain-specific polymorphisms and enriches for mutations most likely introduced during UV mutagenesis.

Following variant calling and annotation, a multi-step filtering pipeline was applied to refine the list of candidate mutations (Fig. 8A), including exclusion of low-confidence calls and prioritization of mutations located within or proximal to annotated genes, which reduced the dataset to a small set of candidate *loci*. From this set, three genes with potential relevance to photosynthetic electron transport and cellular metabolism were prioritized (Fig. 8B). Among these candidates, a mutation was identified in the upstream genomic region of a gene encoding ferredoxin-NADP⁺ reductase (FNR). FNR is a key node in photosynthetic electron partitioning, directing electrons toward NADPH formation or alternative sinks, including hydrogen production. The mutation is located approximately 850 bp upstream of the annotated start codon, within an intergenic region that may function as a regulatory promoter, raising the possibility of an effect on gene expression rather than protein structure^50^.

**Figure 8.**
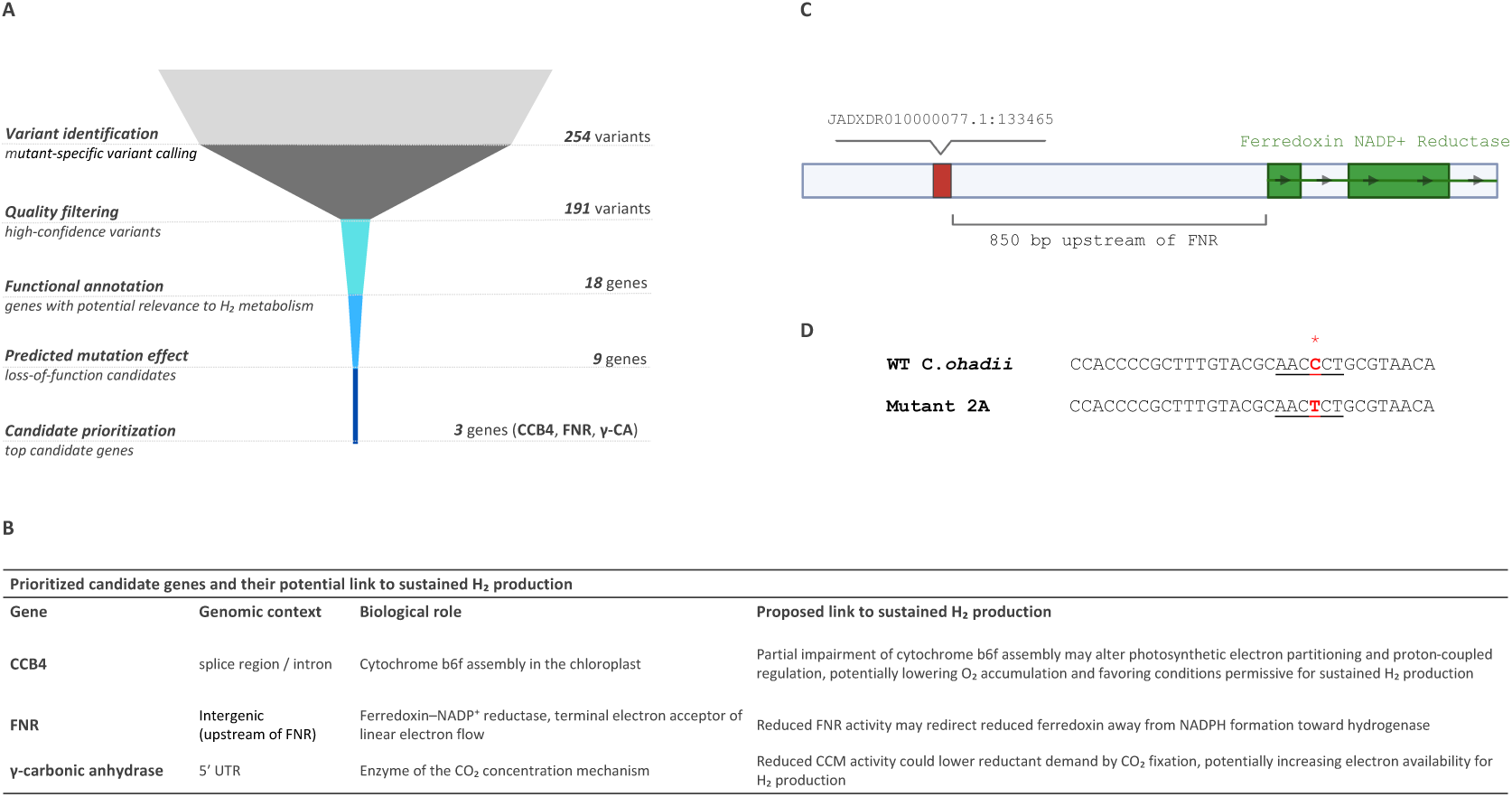
Variant prioritization and validation of candidate loci in the *Chlorella ohadii* mutant 2A. (**A**) Filtering pipeline used to identify candidate variants from whole-genome sequencing of the mutant relative to the parental wild-type (WT) strain. Sequential filtering steps reduced the initial set of variants to three prioritized candidate genes. (**B**) Summary of the prioritized candidate genes, including variant location and biological function. (**C**) Genomic context of the mutation identified upstream of the ferredoxin-NADP^+^ reductase (FNR) gene. The mutation is located approximately 850 bp upstream of the coding region. (**D**) Alignment of the corresponding genomic region in the wild-type strain (*C. ohadii*) and the mutant strain (2A), confirming the nucleotide substitution identified by whole-genome sequencing.

To validate the variant identified by whole-genome sequencing, the corresponding genomic region was amplified and sequenced from both mutant and wild-type strains. Sanger sequencing confirmed the presence of the mutation in the mutant and its absence in the wild type, supporting the accuracy of the variant-calling pipeline (Fig. 8C-D).

This step extends the discovery framework from phenotype identification to candidate gene prioritization, reinforcing the integration of physiological and genetic layers within the platform.

## 4. Discussion

A central challenge in photobiological hydrogen production is not only sustaining hydrogen evolution but doing so within physiological states compatible with biomass retention and industrial deployment. Photobiological hydrogen production in microalgae remains largely confined to laboratory systems, with strain improvement predominantly relying on targeted genetic engineering in a limited number of model organisms. While these approaches provide valuable mechanistic insight, they also generate transgenic strains that face substantial regulatory barriers when algal biomass is intended for food or feed applications. To address this, we demonstrate a non-GMO discovery platform that combines UV mutagenesis with targeted physiological screening to identify hydrogen-producing mutants across multiple algal species. Productive mutants were isolated in three phylogenetically distinct backgrounds, including the industrial strain *Chlorella vulgaris* and the extremophile *Chlorella ohadii*. In addition to enabling cross-species discovery, this platform achieves high screening efficiency, with substantial enrichment at the primary selection stage and markedly improved recovery of hydrogen-producing phenotypes relative to previous approaches.

Despite their diverse genetic backgrounds, all validated mutants converged on a common physiological outcome characterized by limited oxygen accumulation and reduced carbon fixation. However, this state emerged through distinct photosynthetic configurations rather than a single underlying mechanism. Comparative physiological analyses revealed multiple routes toward sustained hydrogen production, including enhanced downstream electron withdrawal coupled with oxygen consumption, reduced PSII electron supply, and condition-dependent redistribution of PSI redox states (Supplementary Table S2). These observations indicate that hydrogen production represents a broadly accessible physiological state of the photosynthetic apparatus that can be reached through multiple perturbations of electron transport and energy balance.

The *Chlorella ohadii* mutant 2A provides a clear example of one such configuration. This strain maintains functional photochemistry and sustained light-dependent O_2_ evolution while preventing net O_2_ accumulation through enhanced downstream consumption. Reduced acceptor-side limitation at PSI (Fig. 7A) supports continued electron throughput toward hydrogen production, defining a system driven by efficient downstream electron withdrawal and active O_2_ turnover. To begin linking these physiological properties to their genetic basis, whole-genome sequencing was performed on mutant 2A. This analysis identified a small set of candidate loci, including a mutation located upstream of a gene encoding ferredoxin-NADP^+^ reductase (FNR), a central regulator of photosynthetic electron partitioning. Although causality remains to be established, these results demonstrate how phenotype-driven discovery can be extended toward genetic hypothesis generation and target prioritization.

Taken together, our findings suggest that sustained hydrogen production is not a rare or highly specialized phenotype but rather one manifestation of a broader photosynthetic design space. The convergence of distinct mutants toward related PSI redox states indicates that hydrogen production is supported by a common downstream electron-limited condition that can be achieved through multiple upstream physiological configurations. While the present study focused primarily on photosynthetic electron transport, additional variation in carbon metabolism and regulatory networks is likely to further expand the range of states compatible with hydrogen production.

Importantly, physiological behaviors observed at laboratory scale must ultimately be evaluated under the constraints of industrial cultivation. While the mutants identified here demonstrate sustained hydrogen production under controlled conditions, future studies will be required to determine whether these traits remain advantageous under large-scale photobioreactor operation. Nevertheless, the framework presented here provides a scalable approach for identifying productive hydrogen-producing phenotypes across diverse algal species. Moreover, mutations identified through phenotype-driven screening define candidate targets for validation and potential transfer into industrial strains using targeted genome-editing approaches that comply with application-specific regulatory frameworks. More broadly, this work demonstrates how systematic exploration of physiological diversity can expand the range of accessible solutions for photobiological hydrogen production and accelerate the development of strains suitable for future deployment.

## 5. Methods

### 5.1 Algal strains and culture conditions

*Chlamydomonas reinhardtii* (CC125 and T222), *Chlorella ohadii*, and *Chlorella vulgaris* were used in this study. The *Chlorella vulgaris* strain was obtained from Bar-Algae (Israel), while all other strains correspond to standard laboratory strains commonly used in algal research. All experiments were conducted in Tris-acetate-phosphate (TAP) medium, supporting mixotrophic growth. Cultures were maintained at 24 °C under continuous illumination at low light intensity (15–30 μmol photons m^−2^ s^−1^) using fluorescent light sources. For all downstream physiological assays, cultures were normalized based on chlorophyll content. For *Chlamydomonas* strains, pigments were extracted in 80% (v/v) acetone and quantified as described in ^51^. For *Chlorella* strains, chlorophyll was extracted in 100% methanol and quantified according to ^52^.

### 5.2 UV mutagenesis and mutant library generation

UV mutagenesis was performed using a UV light source integrated within a biological safety cabinet. Cells were exposed at a fixed distance of approximately 40 cm from the UV source. To calibrate mutagenesis conditions, approximately 100 cells from each strain were plated on TAP agar plates and exposed to increasing durations of UV irradiation. Following incubation under standard growth conditions, survival rates were assessed, and exposure times corresponding to ∼10% survival (∼90% lethality) were determined individually for each strain. For mutant library generation, approximately 5,000 cells per strain were plated on TAP agar and exposed to the strain-specific UV conditions identified during calibration. After incubation, surviving colonies (∼10% of the initial population) were allowed to recover and were used to establish mutant libraries of approximately 500 independent colonies per strain for downstream screening.

### 5.3 High-throughput primary screening under high light

Following UV mutagenesis, mutant libraries were screened using a growth-based assay under low and high light conditions. Individual colonies (∼500 per strain) were inoculated into 96-well plates containing TAP medium and grown under low light conditions (15–30 μmol photons m^−2^ s^−1^) to allow biomass accumulation. The screening workflow was designed to be compatible with both manual and automated operation. For automated handling, single colonies were picked using a Pickolo colony picker (SciRobotics) integrated into a Biomek i7 liquid handling system (Beckman Coulter). This platform was used for colony transfer and for spotting cultures onto agar plates. Cultures were subsequently arrayed onto TAP agar plates in duplicate by spotting 5 μL from each well on TAP agar OmniTray plate, in their respectful coordination from the 96-well. The method leads to the growth of the algae colonies as a circular patch, suitable for easily distinguishing changes in phenotype. One plate was maintained under low light conditions (15–30 μmol photons m^−2^ s^−1^) as a growth control, while the second plate was exposed to high light (∼900 μmol photons m^−2^ s^−1^) to impose light stress. For *Chlorella ohadii*, an additional plate was exposed to direct sunlight to provide a more extreme light-stress condition. Sunlight exposure experiments were conducted outdoors in Tel-Aviv, Israel, during the summer months (June–August). After 5–7 days of incubation, colonies that exhibited normal growth under low light but reduced growth or complete growth arrest under high light were classified as screen-positive candidates.

### 5.4 Hydrogen production measurements by membrane inlet mass spectrometry (MIMS)

Hydrogen production and gas-exchange dynamics were measured using membrane inlet mass spectrometry (MIMS), enabling continuous monitoring of dissolved H_2_ (*m*/*z* 2), O_2_ (*m*/*z* 32), and CO_2_ (*m*/*z* 44). Measurements were performed in an optical chamber (ED-101US/MD, Walz) using a quartz cuvette. Prior to analysis, cultures were adjusted to a chlorophyll concentration of approximately 20 μg Chl mL^−1^ in TAP medium supplemented with 50 mM HEPES. Samples were dark-incubated for 2 h to promote oxygen depletion and establish hydrogen-producing conditions. Following dark incubation, samples were illuminated at 370 μmol photons m^−2^ s^−1^ for 16 min, including a final 3-min pulse of saturating light (∼2500 μmol photons m^−2^ s^−1^), followed by a 2-min dark phase. Illumination was provided using a Dual-PAM-100 system (Heinz Walz GmbH, Germany). Signal normalization and calibration were performed using standard procedures as described previously ^53^.

### 5.5 Growth analysis under standard cultivation conditions

Growth curves of wild-type and mutant strains were assessed under standard cultivation conditions. Cultures were grown in TAP medium under continuous illumination at 100 μmol photons m^−2^ s^−1^ and maintained at 24 °C. Cells were inoculated at an initial optical density of OD₇₅₀ = 0.07 and cultivated for 6 days. Optical density at 750 nm (OD₇₅₀) was recorded at 10-min intervals using a Multi-Cultivator MC 1000 system (Photon Systems Instruments) to generate growth curves and monitor biomass accumulation.

### 5.6 Oxygen evolution measurements and photosynthesis–irradiance (PI) curve analysis

Oxygen dynamics were measured using an optical oxygen sensor (FireSting, PyroScience) equipped with an OXROB3 probe. Measurements were performed in a temperature-controlled measurement system (ALGi dissolved gas analyzer), which integrates illumination and environmental control, using cell suspensions adjusted to approximately 20 μg Chl mL^−1^ in TAP medium supplemented with 50 mM HEPES. For PI curve analysis, 2 mL of cell suspension was placed in the ALGi measurement vial and dark-adapted for 2 min. Samples were then exposed to a series of increasing light intensities spanning 50 to 2000 μmol photons m^−2^ s^−1^. Each light step was followed by a 1-min dark interval to enable estimation of respiration rates. Net O_2_ evolution was determined from the slope of O_2_ concentration during illumination, while gross O_2_ evolution was calculated by correcting for respiration, defined as the O_2_ consumption rate measured during the preceding dark interval. Rates were normalized to chlorophyll content and calculated by linear regression.

### 5.7 Chlorophyll fluorescence measurements by PAM fluorometry

Chlorophyll fluorescence measurements were performed using a Dual Pulse Amplitude Modulated fluorometer (DUAL-PAM-100; Walz) equipped with DUAL-DR and NIR modules. Measurements were conducted in parallel with MIMS gas-exchange assays under identical experimental conditions, as described in Section 5.4. Light intensity was calibrated using a Walz light detector (model US-SQS/L). Fluorescence parameters, including the effective quantum yield of PSII (Y(II)), electron transport rate (ETR), and non-photochemical quenching (NPQ), were calculated according to standard protocols implemented in the instrument software.

### 5.8 PSI redox partitioning and thylakoid energetic measurements using JTS

PSI redox dynamics and thylakoid membrane energetics were assessed using a Joliot-type spectrophotometer (JTS-150, BioLogic, France), following established protocols ^54^. Electrochromic shift (ECS) measurements were used to monitor changes in the proton motive force (pmf) across the thylakoid membrane, detected as absorbance differences at 520–546 nm. Samples were maintained under continuous stirring in an open vessel under oxic conditions and dark-adapted for 30 min prior to measurement. ECS signals were induced by short actinic light pulses and recorded during brief dark intervals to resolve the kinetics of membrane potential formation and relaxation. For comparison across samples, ECS amplitudes were normalized to the ECS signal corresponding to a single PSI charge separation. This reference signal was determined from the average response to a series of saturating laser flashes recorded in the presence of PSII inhibitors (10 μM DCMU and 1 mM hydroxylamine). P700 redox changes were monitored as absorbance differences at 705–740 nm using pulsed detection light. Cells were adjusted to ∼20 μg Chl mL^−1^ and supplemented with 10% Ficoll to prevent sedimentation. For measurements under anaerobic conditions, samples were dark-incubated for 2 h prior to analysis. Steady-state P700 levels under actinic illumination (Ps), maximal oxidation levels during saturating pulses (Pm), and maximal oxidizable P700 (P′m) in the presence of PSII inhibitors were determined. These parameters were used to calculate PSI quantum yield (Y(I)) and the relative contributions of donor-side (YND) and acceptor-side (YNA) limitations.

### 5.9 Whole-genome sequencing and variant analysis

Whole-genome sequencing was performed at QB3 Genomics (University of California, Berkeley, CA, USA; RRID:SCR_022170). Genomic DNA quality was assessed using a Fragment Analyzer system (Agilent) with a large-fragment assay. DNA was fragmented using a Bioruptor Pico sonication system (Diagenode), and sequencing libraries were prepared using the KAPA HyperPrep kit for DNA (KK8504). Truncated universal stub adapters were ligated to fragmented DNA, which was extended with 6 cycles of PCR using unique dual indexing primers to complete the adapters and enrich the libraries for adapter-ligated fragments. Library quality was checked on a Fragment Analyzer (Agilent) using the HS-NGS assay, and library molarity was measured via quantitative PCR with the KAPA Library Quantification Kit (Roche KK4824) on a BioRad CFX Connect thermal cycler. Libraries were pooled by molarity and sequenced on an Illumina NovaSeq X (25B flowcell) for 2 × 150 cycles, targeting 20 Gb of data per sample. Fastq files were generated and demultiplexed using Illumina BCL Convert v4 with default settings. Raw Illumina reads were trimmed using fastp v1.0.1, removing poly-G tails and bases with a Phred quality score < 20, while retaining reads ≥ 35 bp in length. Trimmed reads were aligned to the *Chlorella ohadii* reference genome (NCBI assembly GCA_025026875.1) using BWA v0.7.19. Variant calling was performed with FreeBayes v1.3.10, assuming haploid ploidy. Alignments with mapping quality < 30 and alleles with base quality < 20 were excluded. Variants located in soft-masked or ambiguous genomic regions in *C. ohadii* were removed from downstream analyses. A variant was considered candidate causative if it had a minimum depth of 10 reads, with ≥ 95% of reads supporting the variant in each mutant sample.

### 5.10 Targeted validation by Sanger sequencing

The genomic region upstream of the FNR gene in *Chlorella ohadii* wild-type and mutant 2A strains was amplified by PCR using Q5 High-Fidelity DNA Polymerase (New England Biolabs). The following primers were used for amplification: forward 5′-CGTCGCCCCTTCTCTTCTTT-3′ and reverse 5′-ATCCCATGCATTGCGCCT-3′. PCR products were purified and subjected to Sanger sequencing, and the resulting sequences were aligned to confirm the presence of the mutation. Alignment of wild-type sequences was performed against the *Chlorella ohadii* reference genome (NCBI assembly GCA_025026875.1). Sequencing data are provided as a Source Data file.

## 6. Acknowledgements

We would like to thank the Yacoby lab members. This study was financed by a grant from the Israel Science foundation 941/22, Ministry of science and Technology (MOST) grant 8395 and and the Council for Higher Education (VATAT) Excellence Center -Energy.

## 7. Author contribution

T.E. conceived the study, designed and performed experiments, analyzed the data, and wrote the manuscript. R.A. generated mutant libraries and carried out primary screening for hydrogen-producing candidates. Y.T. developed the automated platform used for high-throughput screening. J.R.M. assisted with JTS-based measurements. S.J. performed the Sanger-based mutation mapping and validation. A.M. and M.B. performed the genomic analysis and variant identification and contributed to data interpretation. I.Y. conceived the study, supervised the project, contributed to experimental design, wrote the manuscript, and secured funding.

## Notes

### Competing Interest Statement

The authors have declared no competing interest.

