## Supplementary material for "Expanding The Algal Hydrogen Toolbox: A Non-GMO Platform Reveals Multiple Physiological Routes To Sustained Hydrogen Production Across Microalgae": Supp

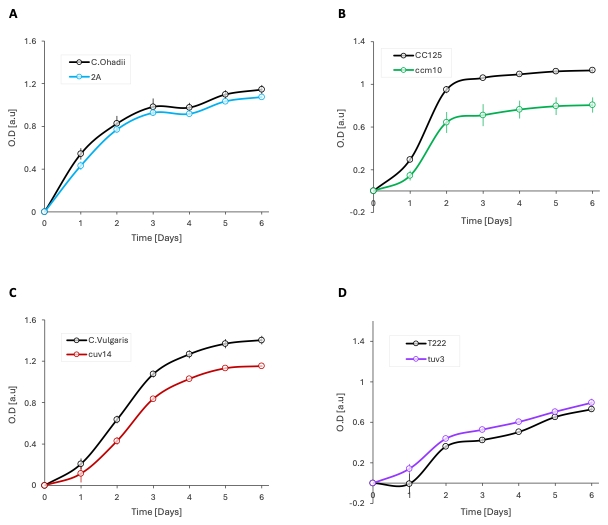

**Figure S1. Growth curves of wild-type and hydrogen-producing mutant strains under mixotrophic conditions.**

Optical density (OD₇₅₀) was monitored over six days in TAP medium under continuous illumination at 100 μmol photons m⁻² s⁻¹. Cultures were inoculated at OD₇₅₀ = 0.07. Each panel compares a wild-type strain (black lines) with its corresponding highest H₂-producing mutant (colored lines). Values represent mean ± SE (n = 4 biological replicates).

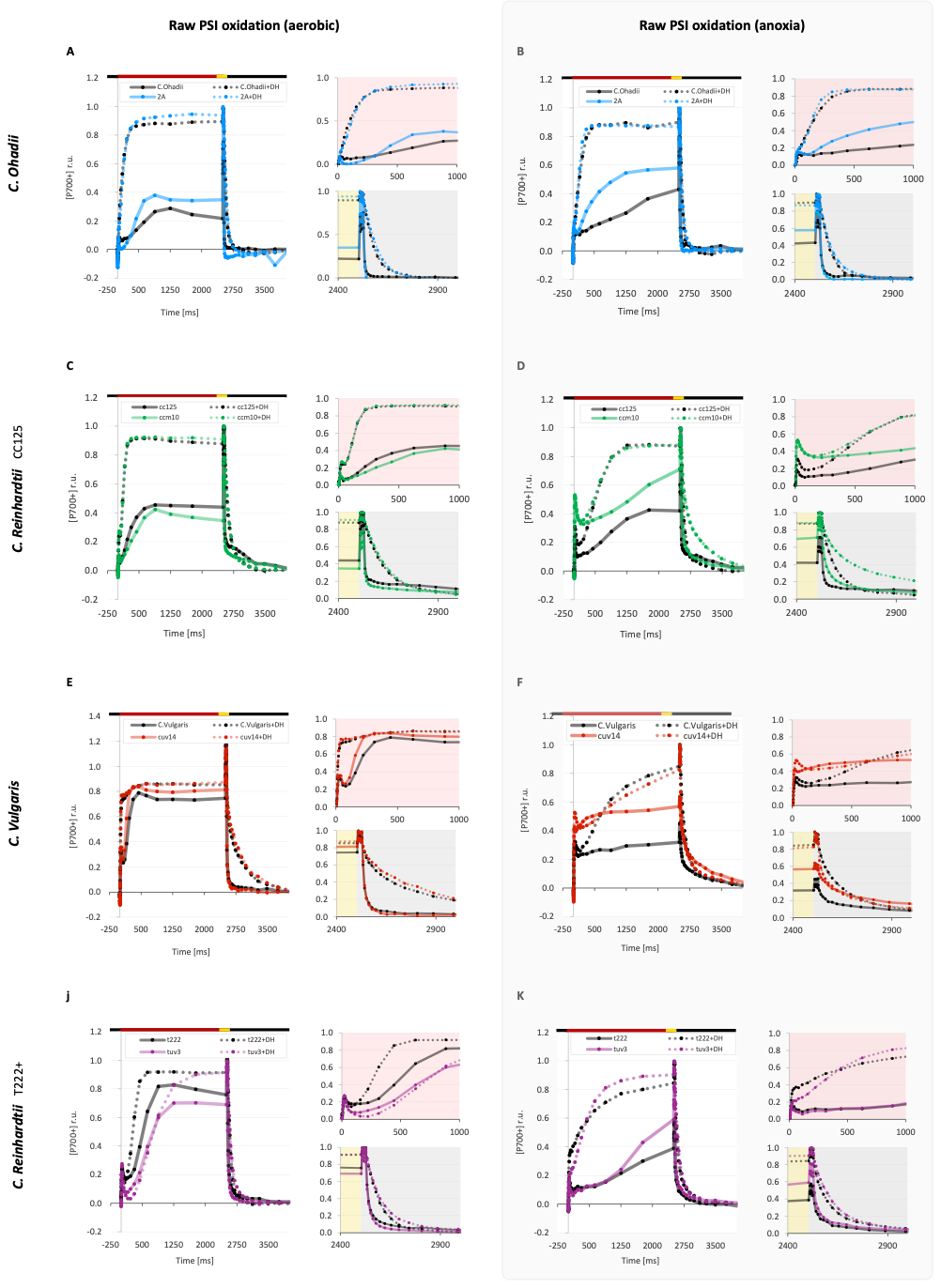

**Figure S2. Representative P700 oxidation kinetics underlying PSI redox partitioning.**

Raw P700⁺ oxidation kinetics recorded using a Joliot-type spectrophotometer (JTS) under aerobic (left column) and anaerobic conditions following 2 h dark incubation (right column). Each row represents one species, shown as wild type (black traces) and its highest hydrogen-producing mutant (colored traces). Illumination regimes are indicated above the traces: black bar, darkness; red bar, actinic light (900 μmol photons m⁻² s⁻¹); yellow bars, saturating pulses (3000 μmol photons m⁻² s⁻¹). Insets show expanded views of the induction and relaxation phases with or without the presence of 10 μM DCMU and 1 mM hydroxylamine (+DH traces) to block PSII.

| Mutant | MIMS signature | O₂ management | PSII (PAM) | PSI redox (P700) | Δψ (ECS) |
| --- | --- | --- | --- | --- | --- |
| **C. ohadii 2A** | Sustained H₂  **↓**low O₂  **↓**CO₂ fixation | High gross O₂ with limited net accumulation | WT-like | ↓ acceptor limitation | WT-like |
| **C.reinhardtii ccm10** |  | Low gross O₂ with restricted net accumulation | ↓ Y(II)  ↓ETR  ↑NPQ | ↑ donor limitation (anoxia)  ↑ acceptor limitation (air) | Slow relaxation |
| **C. vulgaris cuv14** |  | Low gross O₂ with stable gross–net separation | ↓ Y(II)  ↓ETR | ↑ donor limitation (anoxia) | Fast relaxation |
| **C. reinhardtii tuv3** |  | Negative net O₂ at low irradiance; supply-limited balance | ↓ Y(II)  ↓ETR  ↑ NPQ | Condition-dependent redistribution (anoxia vs air) | WT-like |

**Table S2**. **Comparative physiological profiles of hydrogen-producing algal mutants.**

Physiological parameters derived from MIMS (gas exchange), PAM (PSII photochemistry), P700 oxidation (PSI redox partitioning), and ECS (thylakoid membrane potential, Δψ) are summarized relative to the corresponding wild type for each strain. Reported features represent consistent trends observed across measurements. Arrows indicate qualitative changes relative to wild type (increase ↑, decrease ↓), and descriptive terms (e.g., “supply-limited”, “stable gross–net separation”) summarize dominant physiological behaviors inferred from the data.
